# Increased nitrogen fixation and remobilization may contribute to higher seed protein without a yield penalty in a soybean introgression line

**DOI:** 10.1101/602748

**Authors:** Anna M. Locke, Martha E. Ramirez

## Abstract

The development of soybean varieties with higher seed protein concentration has been hindered by a negative correlation between seed protein concentration and yield. Benning HP, a genotype that breaks this tradeoff, contains a high protein allele introgressed into the cultivar Benning. Because seed protein is thought to be limited by N, field and growth chamber experiments were performed to identify the N flux(es) that enable Benning HP’s increased seed protein without a yield penalty. When the N source was completely controlled in growth chambers, Benning HP was able to fix more N than its recurrent parent, but this response depended on the strain of the rhizobia symbiont and was not observed at all developmental stages. In the field, Benning HP remobilized N from its leaves at a higher rate during seed fill, but this response was only observed in one of the years studied. These results demonstrate that Benning HP has higher potential for N fixation and N remobilization from vegetative tissue compared to its lower protein recurrent parent, but those traits are not consistently expressed and may depend on environmental and sink control.

## 1. Introduction

A substantial portion of the value in soybean [*Glycine max* (L.) Merr.] seed is derived from its meal protein, which is an important source of nutrition for poultry and livestock worldwide. Soybean seed protein concentration ranges from 35% to over 50% among genotypes (Hwang et al. 2014, Zhang et al 2018). However, the negative correlation between seed protein concentration and yield is a major hurdle in exploiting this phenotypic diversity to breed agronomically elite, high protein soybean varieties (Brzostowski et al., 2017). While yield has steadily increased, seed protein concentration has decreased with year of cultivar release (Mahmoud et al., 2006; Morrison et al., 2000). It has been hypothesized that the persistence of greenness, which prolongs CO_2_ assimilation during seed fill—increasing yield—has reduced the N available for remobilization from vegetative tissue (Kumudini et al., 2002).

N is primarily transported into the seed coat in the form of ureides and from seed coat to developing seed as glutamine and asparagine (Rainbird et al., 1984b), and energy must be expended in the seed for the synthesis of storage proteins. *In vitro* studies of soybean seed development have found that seed protein accumulation is determined by the supply of N into the seed (Pipolo et al., 2004; Saravitz and Raper, 1995). Similarly, in *Brassica napus* and *Brassica carinata*, final seed protein concentration correlated positively with total amino acid content in leaf phloem sap (Lohaus and Moellers, 2000). When the soybean supply/sink ratio was altered by removing 50% of the pods at each node, seeds accumulated significantly more protein (as % of seed weight) when the assimilate supply per seed was increased (Rotundo et al., 2011).

Three N fluxes can potentially contribute to developing seeds: uptake from the soil, biological N fixation, and remobilization from vegetative tissue. The relative contribution of these components to seed N is variable among genotypes and environmental conditions (Leffel et al., 1992b; Mastrodomenico et al., 2013; Zeiher et al., 1982). It has been hypothesized that the large N demand in developing soybean seeds requires substantial N remobilization from vegetative tissue, triggering leaf senescence and limiting the seed fill period (Sinclair and de Wit, 1976, 1975). Later work found that the proportion of seed N from remobilization to be anywhere from 30% to 100%, depending on genotype (Zeiher et al., 1982). Other studies found that up to 90% of seed N may be supplied by N fixation, and that N fixation continues through most of the seed fill period (Leffel et al., 1992a; Mastrodomenico et al., 2013). A study with three high protein and three average protein soybean genotypes found no correlation between redistributed N from vegetative tissue and total N in mature seeds (Egli and Bruening, 2007). These discrepancies in the estimated contribution of remobilization to seed N may be partly linked to genetic improvement of soybean during the 20^th^ century: when varieties released in the 1930’s were compared with varieties released in the 80’s and 90’s, the earlier genotypes remobilized the same amount of N from vegetative tissue, but the newer genotypes were able to accumulate significantly more N during the seed fill period than the older genotypes (Kumudini et al., 2002).

A genome-wide association study identified 17 loci on 10 chromosomes that are significantly associated with seed protein concentration (Hwang et al., 2014). One of the loci identified in that study, on chromosome 20, resulted in mean protein increase from 41.46% to 44.32%. In a genome-wide analysis including 934 accessions from maturity groups (MG) IV-VI, this high protein locus was refined to a 1 Mbp region (Vaughn et al., 2014), but the specific gene(s) that determine this trait have yet to be resolved. A protein-determining locus in this region of chromosome 20 (hereafter called Gm20) has also been identified in genotypes of *G. max* and *Glycine soja* (Diers et al., 1992; Warrington et al., 2015). One of these analyses was performed in a recombinant inbred population derived from a cross between Benning, a high-yielding MG VII cultivar with moderate seed protein (Boerma et al., 1997), and Danbaekkong (Kim et al., 1996), a lower-yielding MG V cultivar with very high seed protein. This locus explained 55% of the variation in seed protein within the bi-parental population (Warrington et al., 2015). From this population, the Danbaekkong Gm20 allele was backcrossed into Benning to create a near-isogenic line, Benning HP, with ca. 3% increase in seed protein over Benning, similar agronomic traits, and no yield penalty (Prenger et al., 2019).

Benning HP’s seed protein concentration improvement over Benning without yield loss is unique; while the Gm20 allele is linked to seed protein, it does not break through the protein-yield tradeoff in most genetic backgrounds (Brzostowski et al., 2017). An understanding of the physiological process(es) by which Benning HP achieves this increase could lead to new strategies for improving seed protein concentration in other genetic backgrounds. This study aimed to identify differences in N fluxes that contribute to higher seed protein without a yield loss in Benning HP, which possesses the Danbaekkong Gm20 allele, compared to its recurrent parent Benning. N uptake and fixation were evaluated in growth chamber experiments, where N sources and rhizobia could be completely controlled, while N remobilization from aboveground vegetative tissue was evaluated in field experiments.

## 2. Materials and Methods

### 2.1. Field experiment

Seeds of soybean cultivars Benning (Boerma et al., 1997) and Benning HP (Prenger et al., 2019) were sown on 26 May 2016 and 30 June 2017 at Central Crops Research Station in Clayton, NC. Soybeans at this research station were planted in a three-year rotation with corn and cotton. In 2016 the experiment was planted in a Norfolk loamy sand, and in 2017 the experiment was planted in a field with variable Appling sandy loam and eroded Cecil sandy clay loam. Plots were arranged in a randomized complete block design with four replicates of each genotype and 96.5 cm row spacing. Within-row planting density was 39.4 seeds per m. In 2016, plots were 3 rows wide and 3.7 m long; in 2017, plot size was increased to 6 rows wide and 5.5 m long to facilitate destructive aboveground biomass and leaf area index (LAI) measurements.

On field sampling dates, four 2-cm diameter disks were cut from uppermost, fully expanded leaves of two plants per plot on each sampling day, and three pods were removed from main stem nodes in the middle (2016) or upper (2017) third of the canopy from two plants per plot on each sampling day. Samples were dried at 60°C for a minimum of 3 days for C and N analysis.

When Benning and Benning HP were in developmental stage R5 in 2017, all plants in 1 m of a center row in each plot were cut down at the soil level. These rows had not been used for tissue sampling. Plots were harvested at 84 – 88 days after planting (DAP). Leaflets and pods were separated from stems and petioles, and total leaf area was measured with a leaf area meter (LI-3100, LI-COR, Lincoln, NE). All tissues were dried at 60°C for a minimum of 3 days, and leaf, seed, pod, and stem + petiole dry weight was measured for each plant.

At maturity, center rows were harvested with a single-row plot combine (Almaco, Nevada, IA). After seeds were weighed, seed protein concentration was measured using near-infrared spectroscopy (DA-7250, Perten Instruments North America, Springfield, IL). Seed protein concentration is reported on a 13% moisture basis.

### 2.2. Chamber experiment 1

Seeds of Benning and Benning HP were surface sterilized with 95% ethanol and 1% NaOCl (Somasegaran and Hoben, 1985) and incubated in rolled, moistened germination paper at 29°C for two days to germinate. Germinated seeds were transplanted into 0.5 L pots filled with vermiculite. The vermiculite was saturated with water before planting. Three germinated seeds were transplanted per pot and thinned to one plant per pot a week after transplanting. Plants were grown at 26°C day/22°C night with a 9 h photoperiod, with the addition of a 3 h incandescent dark interruption for the first 35 days to suppress flowering.

Five plants per genotype were assigned to each treatment. The four treatments were: (1) NH_4_NO_3_ in nutrient solution, (2) inoculation with *Bradyrhizobium diazoefficiens* strain USDA 110 [formerly classified as *Bradyrhizobium japonicum* (Delamuta et al., 2013)], (3) inoculation with *Bradyrhizobium elkanii* strain USDA 31, or (4) no N fertilizer or inoculation. The rhizobia strains contrast in N fixation efficiency, with *B. diazoefficiens* USDA 110 fixing N more efficiently than *B. japonicum* USDA 31 (Schubert et al., 1978). The N-free plants were included to monitor for potential rhizobia contamination in the substrate and to measure N derived from cotyledons prior to excision. Cotyledons were excised 8 days after transplanting (DAT) to hasten reliance on N treatments. All plants were fed 50% Long-Ashton nutrient solution (Hewitt, 1966), modified by omitting NH_4_NO_3_ for the inoculated plants and the N-free plants and with 7 mM NH_4_NO_3_ for the remaining treatment. Inoculations were conducted immediately after transplanting, by adding 1 mL of the respective rhizobia culture grown in yeast extract-mannitol medium (Somasegaran and Hoben, 1985) and containing approximately 10^8^ CFU ml^-1^ of *B. diazoefficiens* USDA 110 or *B. elkanii* USDA 31 to the transplanted seed. Genotypes and treatments were fully randomized within the growth chamber.

For the first 4 DAT, each pot received 20 – 30 ml of deionized water daily. Beginning at 5 DAT, each pot received 50 ml of water daily, and this was followed by 50 ml of nutrient solution every other day. At 26 DAT, water and nutrient solution were both increased to 80 ml daily. All plants were harvested 45 DAT. Plants were separated into shoots, roots and nodules and dried at 60°C for 3 days for biomass determination, followed by C and N analysis.

### 2.3. Chamber experiment 2

Benning and Benning HP seeds were germinated as described above. Germinated seeds were transplanted into 6 L pots filled with vermiculite and approximately 80 g of crushed oyster shells to control acidification of the rhizosphere in later developmental stages (Israel and Jackson, 1982). The vermiculite was saturated with water before planting. Three germinated seeds were transplanted per pot and thinned to one plant per pot a week after transplanting.

Germinated seeds were inoculated with *B. diazoefficiens* USDA 110 immediately after transplanting as described above and grown in a controlled environment growth chamber under the same conditions as described above. For each genotype, 20 pots were inoculated, and seven pots remained uninoculated. Genotypes and treatments were fully randomized within the growth chamber. For the first four DAT, 30 ml of deionized water were added daily; then, vermiculite was flushed daily with 600 ml of deionized water followed by the addition of 400 ml of N-free nutrient solution. Nutrient solution was modified 50% Long-Ashton solution as described above. After developmental stage V5, water and nutrient solution were increased to 1 L and 400 mL, respectively, twice per day. Cotyledons were excised at 8 DAT.

Five inoculated plants per genotype were harvested at 33, 65, 82, and 132 DAT, when the plants were in developmental stages V5, R5, R6, and R8 (maturity). The non-inoculated plants, which received no N, were also harvested at the first sampling date (33 DAT) to measure N derived from cotyledons prior to excision. At harvest dates, whole plants were separated into nodules, roots, vegetative shoot tissue, and reproductive tissue, and dried at 60°C for a minimum of 3 days. Biomass, C, and N content were measured for dry tissue. Seed protein concentration in mature seed was measured with near-infrared spectroscopy.

### 2.4. Carbon/nitrogen measurements

Samples from the field and from chambers were dried at 60°C for a minimum of 3 days before analysis. Nodules and leaf disks were ground in 2 ml tubes containing two 3-mm stainless steel grinding balls, shaken at 1400 strokes min^-1^ for 45 sec in a homogenizer (Geno/Grinder 2000, Spex CertiPrep, NJ). Whole leaves, stems, seeds, pod shells, and roots were ground using a Wiley mill (Model 4, Thomas Scientific, NJ). Small amounts of seeds, such as the three pod samples collected on field sampling dates, were ground with a centrifugal mill (ZM100, Retsch, Germany). Tissue C and N content was measured with an elemental analyzer (FlashEA 1112, Thermo Scientific, Walham, MA), and the percentage of C and N per sample was calculated with the instrument’s software (Eager Smart, Thermo Scientific, Walham, MA). Biomass and percent N values were multiplied to estimate total N in plant tissues.

### 2.5. Leaf chlorophyll measurements

In the field, two 2-cm diameter disks were cut from uppermost, fully expanded leaves of two plants per plot on each sampling day and flash frozen in liquid N. These disks were sampled from the same leaves on the same days as the disks used for leaf C and N measurements. Leaf disks were stored at −80°C until chlorophyll extraction. Chlorophyll was measured according to Porra et al. (1989). Briefly, frozen leaf disks were ground in chilled methanol. Methanol was decanted and centrifuged at 2500 rpm for 10 min. The supernatant was transferred to a cuvette, and absorbance was measured at 652.0 nm and 665.2 nm. Chlorophyll concentrations in the supernatant (µg/mL) were calculated using the equations:

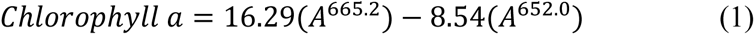

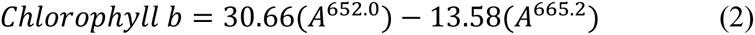

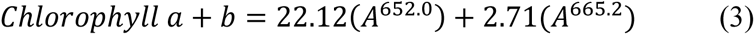

### 2.6. Statistical analysis

Chamber experiments 1 and 2 were completely randomized designs. Two-way analysis of variance (ANOVA) was performed in PROC MIXED (SAS Inc., Cary, NC), and genotype and N treatment (experiment 1) or developmental stage (experiment 2) were treated as fixed effects. Pairwise differences between genotypes within N treatment or developmental stage were calculated with *F*-tests where appropriate using the slice option. *P*-values were adjusted to control the false discovery rate where indicated in tables and figures (Benjamini and Hochberg, 1995).

Each year of the field experiment was a randomized complete block design. The blocking factor was included in all statistical models as a random effect and was dropped from a model when its covariance parameter estimate was equal to or less than 0. Final harvest parameters, LAI, and biomass harvest parameters were analyzed using type 3 ANOVA in PROC MIXED, with genotype treated as a fixed effect and block as a random effect.

To evaluate genotypic differences in leaf and seed N concentration and C/N, seed weight, and leaf chlorophyll concentration throughout each growing season, a linear model with repeated measures was fitted in PROC MIXED with restricted maximum likelihood estimation. Genotype and DAP were fixed effects, block was a random effect, and DAP was the repeated effect with genotype*rep as its subject. Differences between genotypes within DAP were calculated with *F*-tests using the slice option, and the resulting *p*-values were adjusted to control the false discovery rate. To evaluate the rates of change for leaf N concentration, leaf C/N, and leaf chlorophyll during the seed fill period, this model was changed so that DAP was a continuous variable rather than a repeated effect.

In chamber experiment 2 and in the field experiment, biomass parameters and N content were measured at successive developmental stages. These values were used to calculate N lost from vegetative tissue or gained in reproductive tissue between developmental stages. Because total N per organ was destructively measured for different groups of plants at each developmental stage, replicated values for N lost or N gained between stages could not be obtained and tested statistically. However, standard errors (SE) of the means were propagated using the formula:

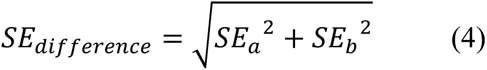

## 3. Results

### 3.1. Benning HP has greater N assimilation potential when inoculated with B. diazoefficiens USDA 110

In chamber experiment 1, plants were supplied with one of three N sources to determine if Benning HP is able to assimilate more N than Benning, and to determine if differences in N assimilation depend on the source of N. The three N sources tested were NH_4_NO_3_ fertilizer, inoculation with *B. diazoefficiens* USDA 110, or inoculation with *B. elkanii* USDA 31. All plants were harvested 45 DAT. Differences in nitrogen supply influenced the rate of development: at 45 DAT, plants inoculated with the more efficient rhizobia strain, USDA 110, were at approximately developmental stage V6, plants inoculated with less efficient rhizobia strain USDA 31 were around V4, and plants fed NH_4_NO_3_ were around V7. Non-inoculated plants did not form nodules, and vermiculite contains no plant-available N, so the N in the no-N plants was derived from stored N prior to cotyledon excision. No-N plants were extremely stunted and chlorotic, as would be expected for severe N deficiency. The average cotyledon N value for each genotype was subtracted from total plant N in the NH_4_NO_3_ fertilizer and rhizobia inoculation treatments to calculate total N assimilated.

N treatment and genotype had a significant effect on total N assimilation, while the genotype × treatment effect was not statistically significant (Table 1). Plants that received NH_4_NO_3_ assimilated the most N per plant (Fig. 1), reflecting the 2-3 weeks after transplanting and inoculation required to develop functional nodules. Plants inoculated with *B. diazoefficiens* USDA 110 fixed more N than plants inoculated with *B. elkanii* USDA 31. Given the low practical risk of accepting a higher type I error rate in the interaction effect to consider the more biologically interesting within-group tests, we conducted *F*-tests for differences between the genotypes within each N treatment. Total N assimilated was not significantly different between genotypes in the plants that were fed fertilizer containing NH_4_NO_3_ (*p* = 0.0828) or inoculated with *B. elkanii* USDA 31 (*p* = 0.5829). When inoculated with *B. diazoefficiens* USDA 110, Benning HP assimilated significantly more N than Benning (*p* = 0.0120) (Fig. 1).

**Figure 1.**
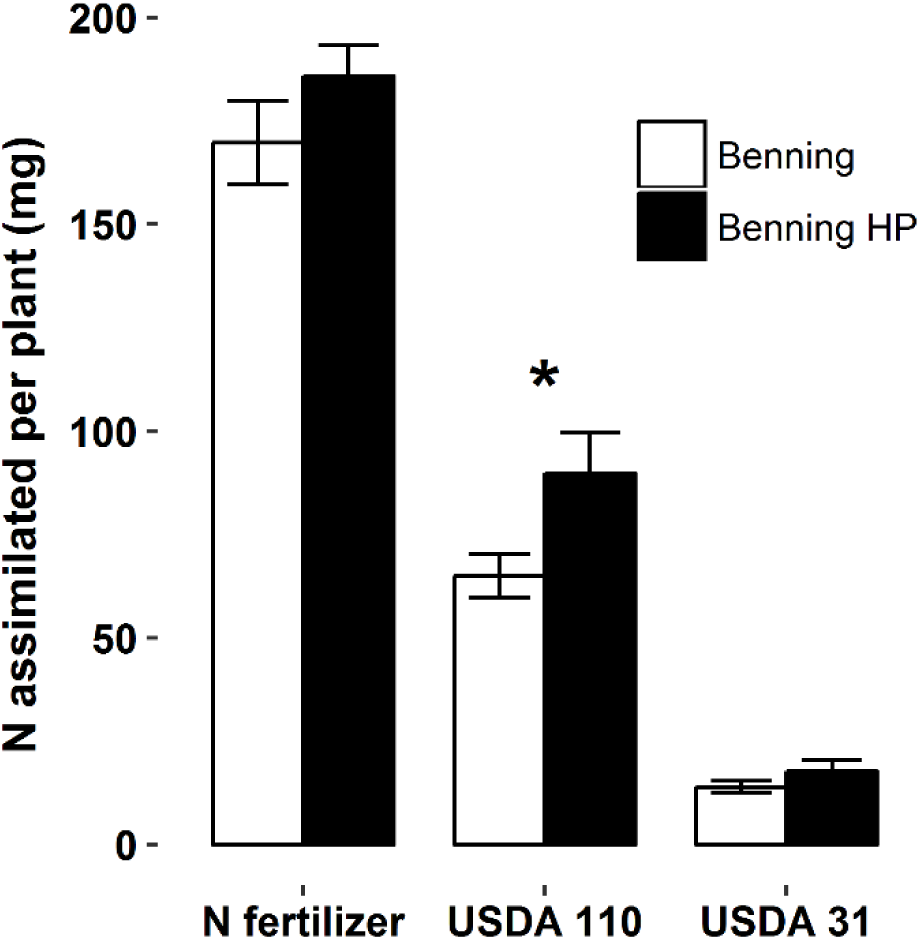
Total nitrogen assimilated per plant (mg) for plants supplied with NH_4_NO_3_ fertilizer, inoculated with *B. diazoefficiens* USDA 110, or inoculated with *B. elkanii* USDA 31 in chamber experiment 1. Whole plants were harvested 45 days after transplanting. Values are mean ± standard error; *n* = 4. Asterisks indicate significant differences (*p* < 0.05) between genotypes within N treatment.

**Table 1.** ANOVA results for chamber experiment 1, comparing N assimilation at 45 DAT in Benning and Benning HP from NH_4_NO_3_ fertilizer, inoculation with *B. diazoefficiens* USDA 110, and *B. elkanii* USDA 31.

| Source | DF | Sum of Squares | Mean Square | Error DF | F Value | *p*-value |
| --- | --- | --- | --- | --- | --- | --- |
| genotype | 1 | 2012.6 | 2012.6 | 30 | 6.87 | **0.0136** |
| N treatment | 2 | 160308 | 80154 | 30 | 273.51 | **< 0.0001** |
| genotype × N treatment | 2 | 663.6 | 331.8 | 30 | 1.13 | 0.3357 |
| residual | 30 | 8791.8 | 293.1 | . | . | . |

Total N assimilated per plant was divided by total nodule weight per plant to calculate N fixed per nodule weight. In chamber experiment 1, Benning HP fixed significantly more N per nodule weight than Benning when inoculated with *B. diazoefficiens* USDA 110 but not when inoculated with *B. elkanii* USDA 31 (Table S1).

### 3.2. Benning HP’s greater N fixation is not observed at every developmental stage

Subsequently, a second growth chamber experiment was conducted to determine if the observed N fixation advantage in Benning HP is consistent throughout development. In chamber experiment 2, Benning and Benning HP were inoculated only with *B. diazoefficiens* USDA 110 and harvested at developmental stages V5, R5, R6, and R8 (maturity). As expected, seed protein concentration at maturity was significantly higher in Benning HP than in its recurrent parent, and seed N concentration and C/N at maturity were also significantly different between the two genotypes (Table 2). Although seed production in a growth chamber does not scale to yield, seed weight per plant at maturity was not significantly different between the two genotypes.

**Table 2.** Seed protein concentration, seed N concentration, seed C/N, and total seed weight per plant at maturity from plants relying on nitrogen fixation with *B. diazoefficiens* USDA 110 in chamber experiment 2. Seed protein concentration is expressed on a 13% moisture basis. *P*-value for genotype effect on each variable shown in bottom row. Values are means for each genotype; *n* = 5.

| Genotype | Seed protein<br>concentration<br>(%) | Seed N<br>concentration<br>(%) | seed C/N | Seed weight<br>per plant (g) |
| --- | --- | --- | --- | --- |
| Benning | 37.5 | 6.5 | 8.2 | 69.0 |
| Benning HP | 41.3 | 7.7 | 6.8 | 82.8 |
| *p*-value | **<0.0001** | **0.0179** | **0.0022** | 0.1554 |

Total N fixed per plant was measured at V5, R5, R6, and R8. Although the developmental stage effect was significant for every organ, the genotype and genotype*stage effects were only significant for whole plants and for reproductive tissue (pods + seeds) (Table 3). Based on pairwise tests within developmental stages, Benning HP had significantly more total N in reproductive tissue only at maturity, which drove the difference in total N fixed at this developmental stage (Fig. 2).

**Figure 2.**
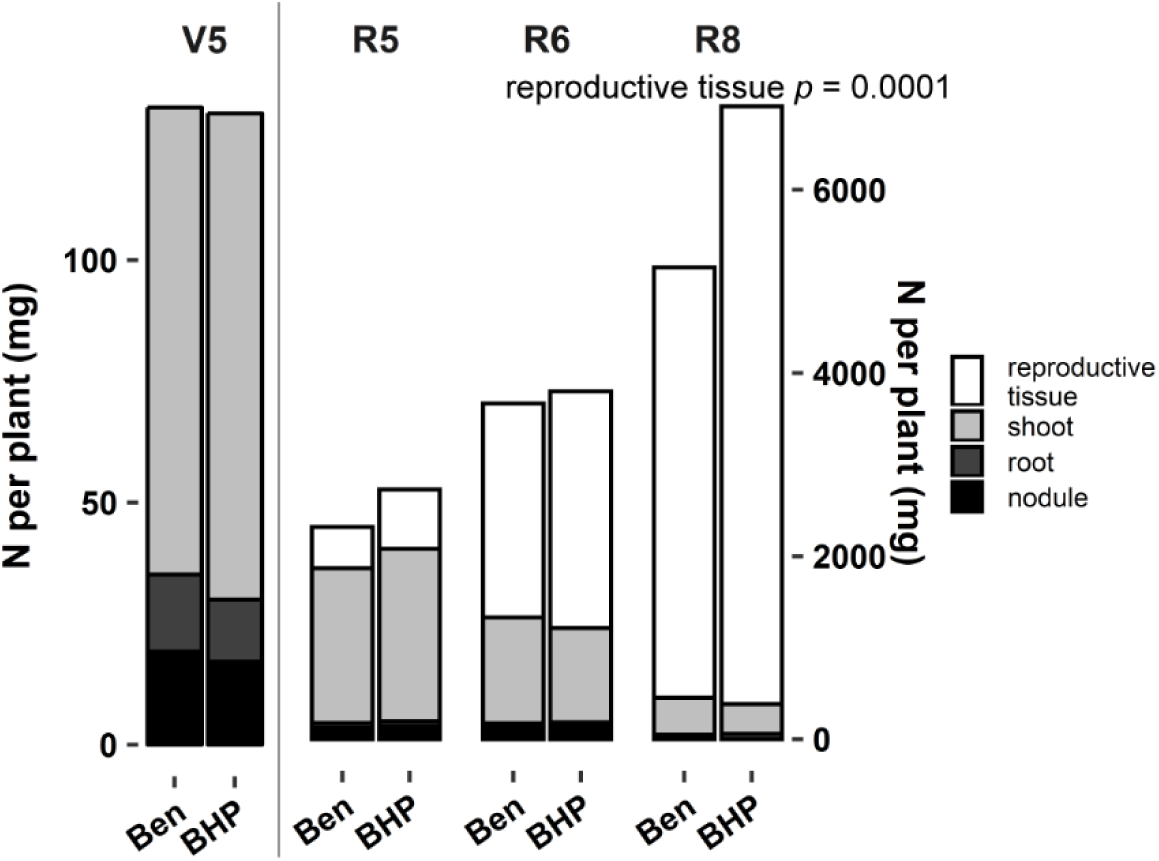
N per plant (mg) in Benning and Benning HP relying on nitrogen fixation with *B. diazoefficiens* USDA 110 in chamber experiment 2. Values are means (*n* = 5). Annotations indicate FDR-corrected *p*-value for significant differences between genotypes in N content per organ; ANOVA results corresponding with these data are presented in Table 3. Scale on left applies to V5; scale on right applies to R5 – R8. Ben = Benning; BHP = Benning HP.

**Table 3.**
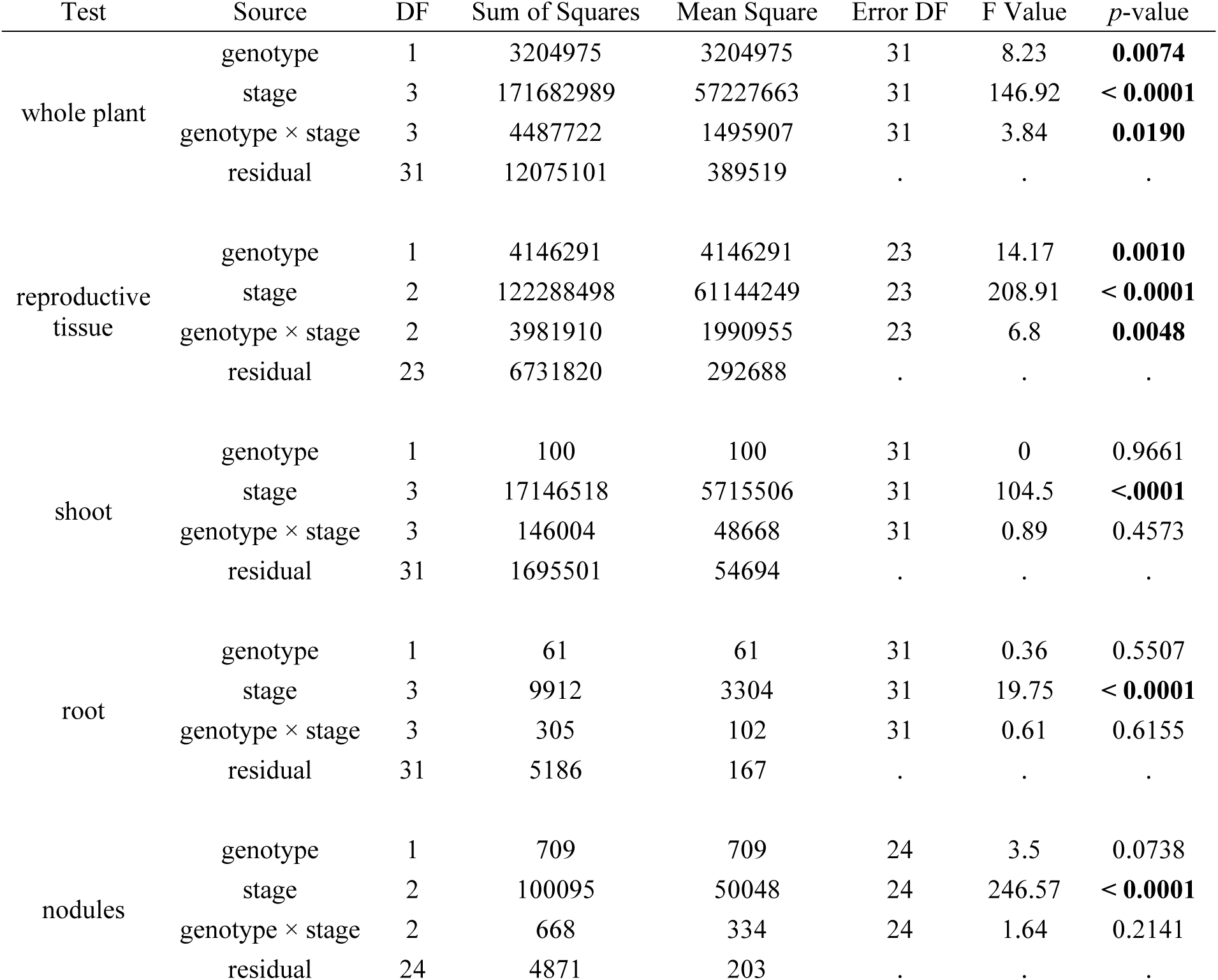
ANOVA results for chamber experiment 2, total nitrogen fixed per plant and nitrogen per organ measured in Benning and Benning HP at four developmental stages. Analysis corresponds with data presented in Figure 2. Reproductive tissue includes seeds and pods.

| Test | Source | DF | Sum of Squares | Mean Square | Error DF | F Value | *p*-value |
| --- | --- | --- | --- | --- | --- | --- | --- |
| whole plant | genotype | 1 | 3204975 | 3204975 | 31 | 8.23 | **0.0074** |
|  | stage | 3 | 171682989 | 57227663 | 31 | 146.92 | **< 0.0001** |
|  | genotype × stage | 3 | 4487722 | 1495907 | 31 | 3.84 | **0.0190** |
|  | residual | 31 | 12075101 | 389519 | . | . | . |
| reproductive tissue | genotype | 1 | 4146291 | 4146291 | 23 | 14.17 | **0.0010** |
|  | stage | 2 | 122288498 | 61144249 | 23 | 208.91 | **< 0.0001** |
|  | genotype × stage | 2 | 3981910 | 1990955 | 23 | 6.8 | **0.0048** |
|  | residual | 23 | 6731820 | 292688 | . | . | . |
| shoot | genotype | 1 | 100 | 100 | 31 | 0 | 0.9661 |
|  | stage | 3 | 17146518 | 5715506 | 31 | 104.5 | **< 0.0001** |
|  | genotype × stage | 3 | 146004 | 48668 | 31 | 0.89 | 0.4573 |
|  | residual | 31 | 1695501 | 54694 | . | . | . |
| root | genotype | 1 | 61 | 61 | 31 | 0.36 | 0.5507 |
|  | stage | 3 | 9912 | 3304 | 31 | 19.75 | **< 0.0001** |
|  | genotype × stage | 3 | 305 | 102 | 31 | 0.61 | 0.6155 |
|  | residual | 31 | 5186 | 167 | . | . | . |
| nodules | genotype | 1 | 709 | 709 | 24 | 3.5 | 0.0738 |
|  | stage | 2 | 100095 | 50048 | 24 | 246.57 | **< 0.0001** |
|  | genotype × stage | 2 | 668 | 334 | 24 | 1.64 | 0.2141 |
|  | residual | 24 | 4871 | 203 | . | . | . |

As in the first chamber experiment, total N assimilated was divided by nodule weight to calculate N fixed per nodule weight. There were not significant differences between genotypes in N fixed per nodule weight at V5, R5, or R6 (Table S1). Insufficient nodule tissue remained for N measurement at R8, so N fixed per nodule weight at maturity could not be calculated.

### 3.3. In growth chambers, Benning HP’s greater reproductive N at maturity must be attributed to N fixation

For chamber experiment 2, the maximum amount of N that could have been remobilized from vegetative tissue into reproductive tissue between developmental stages was estimated from differences in N content between developmental stages. Vegetative N lost was calculated as the difference in vegetative N (leaf, stem, root, and nodule) between developmental stages (Table 4). Because the N remaining in abscised leaves was not measured, this value represents the maximum potential N remobilization from vegetative into reproductive tissue during seed development. Reproductive N gained was calculated as the difference in reproductive N (seed and pod) between developmental stages. In this chamber experiment, the amount of reproductive N gained was always greater than vegetative N lost. Thus, N remobilization cannot have contributed all of the N in reproductive tissue, and the rest of reproductive N must come from N fixation occurring during seed fill. The two genotypes lost similar amounts of N from vegetative tissue and gained similar amounts of N in reproductive tissue from R5 to R6 (Table 4); thus, the estimate for N fixed from R5 to R6 was also similar between the two genotypes. From R6 to R8, the two genotypes again lost similar amounts of N from vegetative tissue. During the same period, however, Benning HP gained substantially more N in its reproductive tissue. Thus, Benning HP may have fixed twice as much N as Benning from R6 to R8, and the propagated standard errors for these estimates do not overlap (Table 4).

**Table 4.** Nitrogen lost per plant (mg) from vegetative tissues and nitrogen gained in reproductive tissues between developmental stages in chamber experiment 2. Nitrogen fixed during each period was estimated as the difference between reproductive N gained and vegetative N lost. Values are mean ± standard error; errors were propagated using equation 4 as described in the Materials and Methods.

| Genotype | R5 – R6 |  |  | R6 – R8 |  |  |
| --- | --- | --- | --- | --- | --- | --- |
|  | Vegetative N lost | Reproductive N gained | Estimated N fixed | Vegetative N lost | Reproductive N gained | Estimated N fixed |
|  | ----- (mg plant<sup>-1</sup>) ----- |  |  |  |  |  |
| Benning | 540 ± 213 | 1887 ± 153 | 1373 ± 262 | 875 ± 164 | 2363 ± 488 | 1488 ± 515 |
| Benning HP | 867 ± 169 | 1940 ± 121 | 1073 ± 208 | 833 ± 46 | 3945 ± 359 | 3112 ± 362 |

### 3.4. Higher N remobilization is not consistent in the field

Benning and Benning HP were grown in the field in 2016 and 2017 to examine nitrogen remobilization during seed fill. Seed protein concentration at harvest was significantly higher in Benning HP than in Benning for both years, while yield was not significantly different between the genotypes (Table 5). Seed N concentration was significantly higher in Benning HP only in 2017, and seed size was significantly smaller in Benning HP in both years.

**Table 5.** Mean yield, seed protein concentration, seed N concentration, and seed size at developmental maturity in field experiments. Seed protein is expressed on a 13% moisture basis; *n* = 4.

| Genotype | 2016 |  |  |  | 2017 |  |  |  |
| --- | --- | --- | --- | --- | --- | --- | --- | --- |
|  | Yield<br>(g m<sup>-2</sup>) | Seed protein<br>concentration<br>(%) | Seed N<br>concentration<br>(%) | Seed size<br>(g 100<br>seed<sup>-1</sup>) | Yield<br>(g m<sup>-2</sup>) | Seed protein<br>concentration<br>(%) | Seed N<br>concentration<br>(%) | Seed size<br>(g 100<br>seed<sup>-1</sup>) |
| Benning | 206.5 | 39.1 | 6.5 | 16.9 | 218.1 | 34.8 | 5.9 | 16.1 |
| Benning<br>HP | 214.6 | 41.9 | 6.8 | 14.8 | 194.3 | 39.0 | 6.5 | 15.1 |
| *p*-value | 0.6005 | **0.0011** | 0.4093 | **0.0406** | 0.1843 | **<0.0001** | **0.0267** | **0.0155** |

LAI and aboveground biomass were measured in 2017 for 1 m rows harvested at DAP 84 – 88. Both genotypes were in developmental stage R5 at this time. LAI and aboveground biomass were not significantly different between genotypes (Table S2). Percent N was measured for aboveground organs to calculate total aboveground N as well as vegetative and reproductive N. There were no differences between the genotypes in total aboveground N, aboveground vegetative N (leaf N + stem N), or reproductive N (seed + pod shell N) at R5 (Table S2). At the time of these measurements, seeds had gained one-third or less of their final dry weight (Fig. 3B).

**Figure 3.**
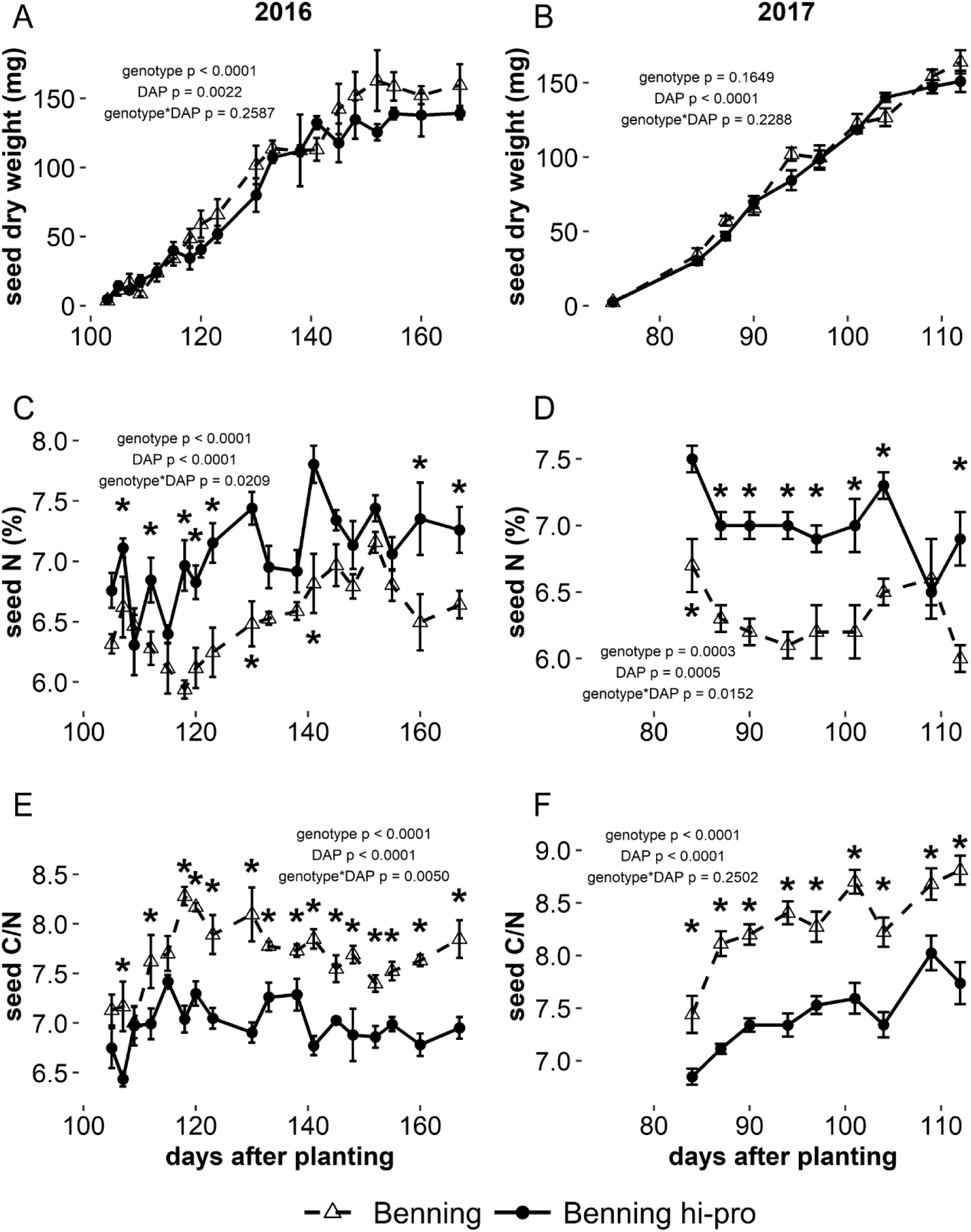
Individual seed weight, seed N concentration, and seed C/N during seed fill for Benning and Benning HP grown in the field. Significant pairwise differences (FDR-corrected *p* < 0.05) between the genotypes within days are indicated by asterisks. Points and error bars are means ± standard error.

The seed N gained from R5 to R8 was calculated as it was for growth chamber experiment 2, from differences in N at each developmental stage (Table S3). For Benning and Benning HP in the field, the amount of seed N gained from R5 to R8 was smaller than the amount of aboveground N in vegetative tissue at R5 which could have been remobilized (Table S2, S3). The standard error estimates for seed N gained overlap greatly. Thus, it was not possible to estimate from these data how much N assimilation occurred during this time period or if N assimilation may have differed between genotypes.

Seed growth in Benning and Benning HP was examined in both years. Seed fill began fewer DAP in 2017, because the planting date was much later that year. In 2016, the overall effect of genotype on individual seed dry weight was significant (Fig. 3A). Individual seed weight tended to be higher in Benning late in seed fill in 2016, but pairwise comparisons were not significant on any specific sampling dates during seed fill, and genotype did not have a significant effect on individual seed weight during seed fill 2017 (Fig. 3B). Genotype had a significant effect on seed N concentration (Fig. 3C – D) and on seed C/N (Fig. 3E – F) throughout seed fill in both years, and the pairwise difference between genotypes was significantly different on almost every measurement day through both years. In 2017, seeds were too small on the first sampling date to measure C and N content.

In leaves, genotype had a significant effect on N concentration and on C/N across the growing season in 2016, but not in 2017 (Fig. S1A, S1B). Differences between the two genotypes were not significant on individual measurement days during vegetative development or during flowering, but leaf N concentration became significantly lower in Benning HP by the end of the seed fill period in 2016 (Fig. S1A), and leaf C/N diverged a few sampling dates earlier (Fig. S1C), suggesting more N depletion from leaves during the seed fill period. The same leaf N concentration and C/N data from the seed fill period were then analyzed with DAP treated as a continuous variable instead of a repeated effect to test if the slope of leaf N concentration and leaf C/N over time was different between the two genotypes. In 2016, genotype and genotype*DAP effects were significant across the seed fill period for leaf C/N and leaf N concentration (Fig. 4A, 4C). The slope of predicted leaf N concentration over time was significantly lower for Benning HP than for Benning, while the slope of predicted leaf C/N over time was significantly higher in Benning HP. However, genotype and genotype*DAP effects were not significantly different in 2017 (Fig. 4B, 4D).

**Figure 4.**
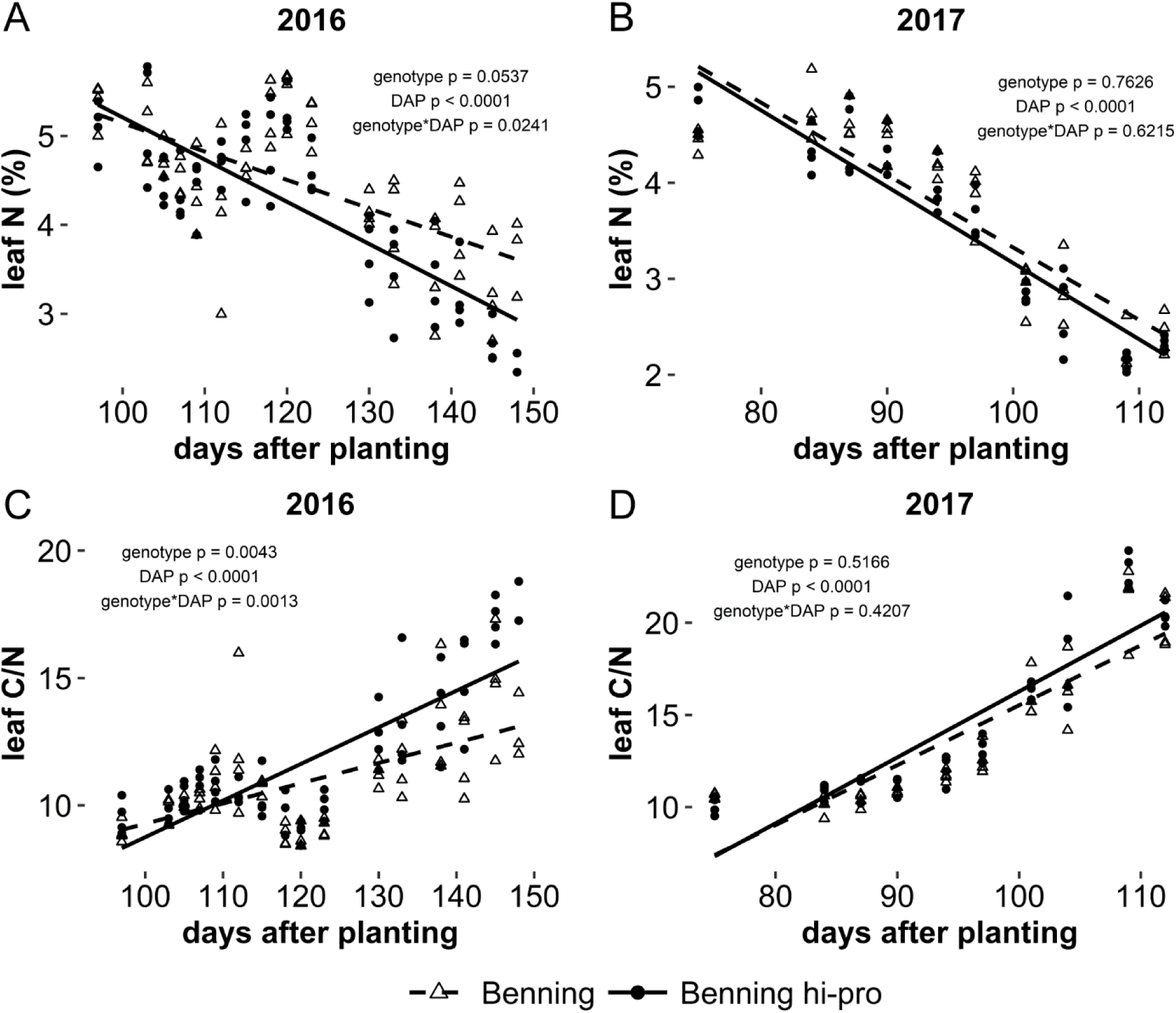
Leaf N concentration and C/N change during the seed fill period, R5 and R6. Dashed and solid lines are predicted values for leaf N concentration and C/N in Benning and Benning HP from the linear model. *P*-values indicate significance of fixed effects in the model.

Leaf chlorophyll was measured for the same DAP to assess whether differences in N remobilization rates could be detected through changes in leaf chlorophyll content. Despite the differences found for leaf N concentration and leaf C/N, the genotype and genotype*DAP effects for leaf chlorophyll in 2016 were not significantly different between Benning and Benning HP (Fig. S2).

## 4. Discussion

The primary goal of this study was to identify the N flux(es) that enable higher seed protein without a yield penalty in Benning HP compared to its recurrent parent, Benning. In these experiments, Benning HP was capable of greater N fixation when N sources were controlled in growth chambers and of a higher rate of N depletion from vegetative tissue during seed fill in the field, but neither of these traits was observed consistently. Strikingly, Benning HP plants inoculated with *B. diazoefficiens* USDA 110 in growth chambers fixed over twice as much N on average from R6 to maturity. Differences in seed protein concentration between two unrelated genotypes of similar yield potential have previously been linked to higher N fixation during seed fill period (Leffel et al., 1992a). In a greenhouse study, N fixation across reproductive development, measured by integrating acetylene reduction measurements over time, was found to be significantly and positively correlated with seed protein concentration, while integrated N fixation was not significantly correlated with yield (Fabre and Planchon, 2000).

*B. diazoefficiens* USDA 110 is more efficient at fixing N than *B. elkanii* USDA 31, and this has been linked to the presence of a hydrogenase in the nodule which recycles H_2_ evolved from the nitrogenase-catalyzed reaction (Caldwell and Vest, 1970; Schubert et al., 1978). As expected, plants inoculated with *B. diazoefficiens* USDA 110 in this study fixed more N and had greater dry weight across genotypes than those inoculated with *B. elkanii* USDA 31. The lower N fixation efficiency of *B. elkanii* USDA 31 restricted the higher N fixation potential of Benning HP that was observed with *B. diazoefficiens* USDA 110. The native rhizobia population was not characterized for the field experiment, but a prior analysis in a different soil type at the same research station found the majority of nodules to be infected with rhizobia belonging to serotypes 31/94, 46/76, and 76 (Ramirez et al., 1997). In that study, only about 15% of the samples isolated from the research station had high N fixation capacity, as does *B. diazoefficiens* USDA 110, while ca. 20% had low N fixation capacity like *B. elkanii* USDA 31, and the rest had intermediate N fixation capacity. Thus, it is possible that in our field experiment, Benning HP’s potential to supply more N to developing seeds from N fixation was restricted by the efficiency of rhizobia strain(s) that were present in soil.

The rate of N remobilization, measured as the slope of leaf C/N over time, from Benning HP leaves was only higher in one of the two years of the field experiment, despite Benning HP maintaining higher seed protein concentration and similar yield to Benning in both years. This variability in N remobilization between years, together with the growth chamber results, do not support the self-destructive hypothesis, which suggested that high seed N requirements in soybean trigger leaf senescence and thus limit the seed fill period (Sinclair and de Wit, 1975). Under the artificial, but optimal, environmental conditions of the growth chambers, the high N requirement in developing seeds was met by sustained N fixation though seed fill in both Benning and Benning HP, as remobilized N accounted for less than half of the N in each genotype at maturity. In the field, Benning HP produced the same seed yield as Benning despite remobilizing N from leaves at a higher rate in 2016. This suggests that under favorable environmental conditions, leaves can continue to assimilate carbon and thus fuel N fixation well into the seed fill period, and that environmental cues may play a larger role than seed N demand in triggering leaf senescence. Because the carbon demand for N fixation is high (Rainbird et al., 1984a), this implies that leaf senescence may actually have been delayed relative to seed development if photosynthesis continued at rates necessary to sustain nodule function.

The data from these experiments cannot conclusively determine if seed composition differences in Benning HP are source- or sink-driven. However, differences in seed N concentration and seed C/N between Benning and Benning HP were established early in seed development, when seeds had gained less than a third of their final weight. Leaf C/N had not yet diverged between the two genotypes this early in seed development. These observations, together with the intermittent differences in N fixation and N remobilization rate, suggest that the Benning HP’s higher seed N may be more dependent on N sink strength than on uniformly higher N fixation or N remobilization.

In growth chamber experiment 2, the higher total N assimilation in Benning HP at maturity was accompanied by substantially higher mature seed weight per plant (ca. 20%), congruent with a previous finding that total N at R7 is positively correlated with yield across genotypes (Rotundo et al., 2014). The seed protein gain for chamber-grown Benning HP was similar to that in the field, ca. 3-4%. As Benning HP’s maximum protein gain over Benning seems to be in this 3-4% range, the potential for greater N fixation during seed fill in Benning HP could contribute to yield maintenance despite higher seed protein.

In the field, individual seed weight was lower in Benning HP when considered across the seed fill period in 2016, and Benning HP’s 100 seed weight at harvest was significantly lower in both years. This is consistent with the findings of Prenger et al. (2019). Based on these findings, it is possible that Benning HP’s higher seed protein concentration is related to reduced weight accumulation per seed relative to N, but yield is maintained because the of Benning HP’s ability to remobilize or fix sufficient N to support a similar total weight of seeds. Genotypic differences in seed sink strength for N were also observed in an *in vitro* study comparing cotyledon N uptake by normal and high protein genotypes at a range of N concentrations (Hayati et al., 1996). In that study, cotyledon N concentration in both genotypes responded positively to N availability in solution, but the high protein genotype’s seeds were able to take up more N at every concentration. Seed N concentration was significantly different between Benning and Benning HP through most of seed development. The lack of significance for the seed N concentration in 2016 harvest data is probably the result of variability through the canopy (Huber et al., 2016). Harvest samples were taken from the combine, which homogenized all the seed from a row. In contrast, samples during seed development were selected from the same canopy position.

Directly measuring leaf N requires specialized, expensive instrumentation, and requires destructive sampling. Chlorophyll content also declines as leaves senesce, and chlorophyll can be extracted in relatively simple assay (Porra et al., 1989) or estimated non-destructively in the field (e.g., Lichtenthaler et al., 1996; Cassol et al., 2008; Steele et al., 2008). The N in chlorophyll is not directly exported from senescing leaves (Hörtensteiner and Feller, 2002), but N from the concurrent breakdown of chloroplast proteins is an important source of N for remobilization to vegetative tissue (Fischer, 2007; Liu et al., 2008). Thus, the decline in leaf chlorophyll content could be a convenient proxy for estimating N remobilization in future studies. We measured leaf chlorophyll in 2016 to determine if it could be used as a surrogate for N remobilization rates. However, the decline of leaf chlorophyll content during seed fill did not reveal the difference in N depletion that was measured in the field in 2016. These results indicate that chlorophyll would not be reliable for detecting subtle but significant differences in leaf N metabolism.

These experiments revealed that the introgressed region of chromosome 20 from Danbaekkong in the Benning background can increase N fixation as well as the rate of N remobilization from senescing soybean leaves, but that the phenotypic expression of these traits may depend on environment and biotic interactions. Whether the changes in these N fluxes are source or sink driven remains unresolved, and the molecular mechanisms behind these changes have yet to be elucidated. Although the Gm20 protein locus has been narrowed to a <1Mb region (Vaughn et al., 2014), the protein gain combined with maintaining the high yield potential of Benning evidently involves multiple genes, as evidenced by large region of Danbaekkong’s chromosome 20 that is present in Benning HP (Prenger et al., 2019). If the gene(s) involved can be identified, a new path to improving soybean seed composition through breeding or biotechnology could be illuminated.

## Acknowledgement

We thank Thomas E. Carter Jr., Paul Cook, Amy Niewoehner, Heather Sims, Jason Pleasant, Cory Callahan, and John Graeber for assistance with planting, tissue sampling, and harvest. We thank Cathy Herring and Travis Lassiter at Central Crops Research Station for field management. We thank Zenglu Li for generously sharing seed for these studies. This work was supported by USDA-ARS [project 6070-21220-069-00-D].

## Appendix A. Supplementary material

Table S1. N fixed per nodule weight means (mg/g; top) and *p*-values from ANOVA (bottom) in chamber experiments 1 and 2. Letters indicate significant differences between genotypes within a treatment (p < 0.05).

Table S2. Leaf area index, aboveground biomass, and aboveground N measured at 84 – 88 DAP in 2017, when Benning and Benning HP were in developmental stage R5; *n* = 4.

Table S3. Seed N gained from R5 to R8 in the field in 2017. Seed N gained was calculated as the difference in mean seed N between R5 and R8. Values are mean ± standard error; errors were propagated as described in the Methods; *n* = 4.

Figure S1. Leaf N concentration and leaf C/N across the growing season for Benning and Benning HP grown in the field in 2016 and 2017. Leaf tissue was sampled from the uppermost, fully expanded leaf in the canopy. Significant pairwise differences (FDR-corrected *p* < 0.05) between genotypes on a sampling day are indicated by asterisks. Points and error bars are means ± standard error.

Figure S2. Leaf chlorophyll content during the 2016 seed fill period, R5 and R6. Chlorophyll was isolated from tissue sampled from the uppermost, fully expanded leaf in the canopy. Slopes were not significantly different between the two genotypes.

## References

Benjamini, Y., Hochberg, Y., 1995. Controlling the false discovery rate: a practical and powerful approach to multiple testing. J. R. Stat. Soc. Ser. B 57, 289–300.

Boerma, H.R., Hussey, R.S., Phillips, D. V, Wood, E.D., Rowan, G.B., Finnerty, S.L., 1997. Registration of “Benning” soybean. Crop Sci. 37, 1982. https://doi.org/10.2135/cropsci1997.0011183X003700060061x

Brzostowski, L.F., Pruski, T.I., Specht, J.E., Diers, B.W., 2017. Impact of seed protein alleles from three soybean sources on seed composition and agronomic traits. Theor. Appl. Genet. 130, 2315–2326. https://doi.org/10.1007/s00122-017-2961-x

Caldwell, B.E., Vest, G., 1970. Effects of *Rhizobium japonicum* strains on soybean yields. Crop Sci. 19, 19–21.

Cassol, D., De Silva, F.S.P., Falqueto, A.R., Bacarin, M.A., 2008. An evaluation of non-destructive methods to estimate total chlorophyll content. Photosynthetica 46, 634–636. https://doi.org/10.1007/s11099-008-0109-6

Delamuta, J.R.M., Ribeiro, R.A., Ormeño-Orrillo, E., Melo, I.S., Martínez-Romero, E., Hungria, M., 2013. Polyphasic evidence supporting the reclassification of *Bradyrhizobium japonicum* group Ia strains as *Bradyrhizobium diazoefficiens* sp. nov. Int. J. Syst. Evol. Microbiol. 63, 3342–3351. https://doi.org/10.1099/ijs.0.049130-0

Diers, B.W., Fehr, W.R., Shoemaker, R.C., 1992. RFLP analysis of soybean seed protein and oil content. Theor. Appl. Genet. 83, 608–612.

Egli, D.B., Bruening, W.P., 2007. Nitrogen accumulation and redistribution in soybean genotypes with variation in seed protein concentration. Plant Soil 301, 165–172. https://doi.org/10.1007/s11104-007-9434-y

Fabre, F., Planchon, C., 2000. Nitrogen nutrition, yield and protein content in soybean. Plant Sci. 152, 51–58. https://doi.org/10.1016/S0168-9452(99)00221-6

Fischer, A.M., 2007. Nutrient remobilization during leaf senescence, in: Gan, S. (Ed.), Senescence Processes in Plants. Blackwell Publishing Ltd, Oxford, UK, pp. 87–107. https://doi.org/10.1002/9780470988855.ch5

Hayati, R., Egli, D.B., Crafts-Brandner, S.J., 1996. Independence of nitrogen supply and seed growth in soybean: studies using an *in vitro* culture system. J. Exp. Bot. 47, 33–40. https://doi.org/10.1093/jxb/47.1.33

Hewitt, E.J., 1966. Sand and water culture methods used in the study of plant nutrition, 2nd ed. Commonwealth Agricultural Bureau, London.

Hörtensteiner, S., Feller, U., 2002. Nitrogen metabolism and remobilization during senescence. J. Exp. Bot. 53, 927–937. https://doi.org/10.1093/jexbot/53.370.927

Huber, S., Li, K., Nelson, R., Ulyanov, A., DeMuro, C., Baxter, I., 2016. Canopy position has a profound effect on soybean seed composition. PeerJ 4, e2452. https://doi.org/10.7717/peerj.2452

Hwang, E.-Y., Song, Q., Jia, G., Specht, J.E., Hyten, D.L., Costa, J., Cregan, P.B., 2014. A genome-wide association study of seed protein and oil content in soybean. BMC Genomics 15, 1. https://doi.org/10.1186/1471-2164-15-1

Israel, D.W., Jackson, W.A., 1982. Ion balance, uptake, and transport processes in N_2_-fixing and nitrate- and urea-dependent soybean plants. Plant Physiol. 69, 171–178. https://doi.org/10.1104/pp.69.1.171

Kim, S.-D., Hong, E.-H., Kim, Y.-H., Lee, S.-H., Seong, Y.-K., Park, K.-Y., Lee, Y.-H., Hwang, Y.-H., Park, E.-H., Kim, H.-S., Ryu, Y.-H., Park, R.-K., Kim, Y.-S., 1996. A new high protein and good seed quality soybean variety “Danbaegkong.” RDA J. Agric. Sci. 38, 228–232.

Kumudini, S., Hume, D.J., Chu, G., 2002. Genetic improvement in short-season soybeans: II. Nitrogen accumulation, remobilization, and partitioning. Crop Sci. 42, 141–145.

Leffel, R.C., Cregan, P.B., Bolgiano, A.P., Thibeau, D.J., 1992a. Nitrogen metabolism of normal and high-seed-protein soybean. Crop Sci. 32, 747–750. https://doi.org/10.2135/cropsci1992.0011183X003200030034x

Leffel, R.C., Cregen, P.B., Bolgiano, A.P., 1992b. Nitrogen metabolism of soybean genotypes selected for seed composition, fasciated stem, or harvest index. Crop Sci. 32, 1428–1432.

Lichtenthaler, H.K., Gitelson, A., Lang, M., 1996. Non-destructive determination of chlorophyll content of leaves of a green and an aurea mutant of tobacco by reflectance measurements. J. Plant Physiol. 148, 483–493. https://doi.org/10.1016/S0176-1617(96)80283-5

Liu, J., Yun, H.W., Jun, J.Y., Yu, D.L., Fa, F.S., 2008. Protein degradation and nitrogen remobilization during leaf senescence. J. Plant Biol. 51, 11–19. https://doi.org/10.1007/BF03030735

Lohaus, G., Moellers, C., 2000. Phloem transport of amino acids in two *Brassica napus* L. genotypes and one *B. carinata* genotype in relation to their seed protein content. Planta 211, 833–40. https://doi.org/10.1007/s004250000349

Mahmoud, A.A., Natarajan, S.S., Bennett, J.O., Mawhinney, T.P., Wiebold, W.J., Krishnan, H.B., 2006. Effect of six decades of selective breeding on soybean protein composition and quality: a biochemical and molecular analysis. J. Agric. Food Chem. 54, 3916–3922. https://doi.org/10.1021/jf060391m

Mastrodomenico, A.T., Purcell, L.C., Andy King, C., 2013. The response and recovery of nitrogen fixation activity in soybean to water deficit at different reproductive developmental stages. Environ. Exp. Bot. 85, 16–21. https://doi.org/10.1016/j.envexpbot.2012.07.006

Morrison, M.J., Voldeng, H.D., Cober, E.R., 2000. Agronomic Changes from 58 Years of Genetic Improvement of Short-Season Soybean Cultivars in Canada. Agron. J. 784, 780–784.

Pipolo, A.E., Sinclair, T.R., Camara, G.M.S., 2004. Protein and oil concentration of soybean seed cultured in vitro using nutrient solutions of differing glutamine concentration. Ann. Appl. Biol. 144, 223–227. https://doi.org/10.1111/j.1744-7348.2004.tb00337.x

Porra, R.J., Thompson, W.A., Kriedemann, P.E., 1989. Determination of accurate extinction coefficients and simultaneous equations for assaying chlorophylls a and b extracted with four different solvents : verification of the concentration of chlorophyll standards by atomic absorption spectroscopy. Biochim. Biophys. Acta 975, 384–394.

Prenger, E.M., Ostezan, A., Mian, R., Buckley, B., Stupar, R.M., Glenn, T., Li, Z., 2019. Introgression of a high protein allele into an elite soybean variety results in a high-protein near-isogenic line with yield parity. Crop Sci. 59, 1–11. https://doi.org/10.2135/cropsci2018.12.0767

Rainbird, R.M., Hitz, W.D., Hardy, R.W., 1984a. Experimental determination of the respiration associated with soybean/rhizobium nitrogenase function, nodule maintenance, and total nodule nitrogen fixation. Plant Physiol. 75, 49–53. https://doi.org/10.1104/pp.75.1.49

Rainbird, R.M., Thorne, J.H., Hardy, R.W., 1984b. Role of amides, amino acids, and ureides in the nutrition of developing soybean seeds. Plant Physiol. 74, 329–334.

Ramirez, M.E., Israel, D.W., Wollum II, A.G., 1997. Phenotypic characterization of soybean Bradyrhizobia in two soils of North Carolina. Soil Biol. Biochem. 29, 1547–1555.

Rotundo, J.L., Borrás, L., de Bruin, J.D., Pedersen, P., 2014. Soybean nitrogen uptake and utilization in Argentina and United States cultivars. Crop Sci. 54, 1153–1165. https://doi.org/10.2135/cropsci2013.09.0618

Rotundo, J.L., Borrás, L., Westgate, M.E., 2011. Linking assimilate supply and seed developmental processes that determine soybean seed composition. Eur. J. Agron. 35, 184–191. https://doi.org/10.1016/j.eja.2011.05.002

Saravitz, C.H., Raper, C.D., 1995. Responses to sucrose and glutamine by soybean embryos grown in vitro. Physiol. Plant. 93, 799–805. https://doi.org/10.1111/j.1399-3054.1995.tb05134.x

Schubert, K.R., Jennings, N.T., Evans, H.J., 1978. Hydrogen reactions of nodulated leguminous plants: II. Effects on dry matter accumulation and nitrogen fixation. Plant Physiol. 61, 398– 401. https://doi.org/10.1104/pp.61.3.398

Sinclair, T.R., de Wit, C.T., 1976. Analysis of the carbon and nitrogen limitations to soybean yield. Agron. J. 68, 319–324.

Sinclair, T.R., de Wit, C.T., 1975. Photosynthate and nitrogen requirements for seed production by various crops. Science 189, 565–567. https://doi.org/10.1126/science.189.4202.565

Somasegaran, P., Hoben, H.J., 1985. Methods in Legume-Rhizobium Technology. University of Hawaii Department of Agronomy and Soil Science, Paia, HI.

Steele, M., Gitelson, A.A., Rundquist, D., 2008. Nondestructive estimation of leaf chlorophyll content in grapes. Am. J. Enol. Vitic. 59, 299–305. https://doi.org/10.2307/2445170.

Vaughn, J.N., Nelson, R.L., Song, Q., Cregan, P.B., Li, Z., 2014. The genetic architecture of seed composition in soybean is refined by genome-wide association scans across multiple populations. G3 4, 2283–2294. https://doi.org/10.1534/g3.114.013433

Warrington, C. V., Abdel-Haleem, H., Hyten, D.L., Cregan, P.B., Orf, J.H., Killam, A.S., Bajjalieh, N., Li, Z., Boerma, H.R., 2015. QTL for seed protein and amino acids in the Benning × Danbaekkong soybean population. Theor. Appl. Genet. 128, 839–850. https://doi.org/10.1007/s00122-015-2474-4

Zeiher, C., Egli, D.B., Leggett, J.E., Reicosky, D.A., 1982. Cultivar differences in N redistribution in soybeans. Agron. J. 74, 375–379.

