## Supplementary Material for "Increased nitrogen fixation and remobilization may contribute to higher seed protein without a yield penalty in a soybean introgression line"

Table S1. N fixed per nodule weight means (mg/g; top) and *p*-values from ANOVA (bottom) in chamber experiments 1 and 2. Letters indicate significant differences between genotypes within a treatment ($p < 0.05$).

| | Chamber Experiment 1 | | | Chamber Experiment 2 | | |
|---|---|---|---|---|---|---|
| | *Bradyrhizobia* strain | | | Developmental Stage | | |
| Genotype | *B. elkanii* USDA 31 | *B. diazoefficiens* USDA 110 | | V5 | R5 | R6 |
| Benning | 284.7 | 406.3 **a** | | 486.51 | 1368.62 | 1877.51 |
| Benning hi-pro | 267.1 | 504.6 **b** | | 509.39 | 1374.95 | 1899.56 |
| Source of variation | | | Source of variation | | | |
| Genotype (G) | 0.1647 | | Genotype (G) | | 0.7379 | |
| Treatment (T) | **<0.0001** | | Stage (S) | | **<0.0001** | |
| G × T | 0.0512 | | G × S | | 0.9887 | |

Table S2. Leaf area index, aboveground biomass, and aboveground N measured at 84 – 88 DAP in 2017, when Benning and Benning HP were in developmental stage R5; *n* = 4.

| Genotype | Leaf area index | Aboveground biomass | Aboveground N | Vegetative N | Reproductive N |
|---|---|---|---|---|---|
| | | ———— ($g\ m^{-2}$) ———— | | | |
| Benning | 4.484 | 638.2 | 19.9 | 11.2 | 8.6 |
| Benning HP | 4.359 | 595.7 | 18.2 | 9.8 | 8.4 |
| *p*-value | 0.8097 | 0.5709 | 0.4713 | 0.3587 | 0.8660 |

Table S3. Seed N gained from R5 to R8 in the field in 2017. Seed N gained was calculated as the difference in mean seed N between R5 and R8. Values are mean ± standard error; errors were propagated as described in the Methods; *n* = 4.

| Genotype | Seed N at R5 | Seed N at R8 | Seed N gained R5 – R8 | Vegetative N at R5† |
|---|---|---|---|---|
| | ——————— g $m^{-2}$ ——————— | | | |
| Benning | 5.9 ± 0.5 | 12.9 ± 1.1 | 7.0 ± 1.2 | 11.2 |
| Benning hi-pro | 6.2 ± 1.0 | 12.7 ± 0.9 | 6.5 ± 1.3 | 9.8 |
| *p*-value | 0.8240 | 0.8473 | — | 0.3587 |

† Vegetative N data from Table S2, shown again here for ease of comparison

Figure S1. Leaf N concentration and leaf C/N across the growing season for Benning and Benning HP grown in the field in 2016 and 2017. Leaf tissue was sampled from the uppermost, fully expanded leaf in the canopy. Significant pairwise differences (FDR-corrected $p < 0.05$) between genotypes on a sampling day are indicated by asterisks. Points and error bars are means ± standard error.

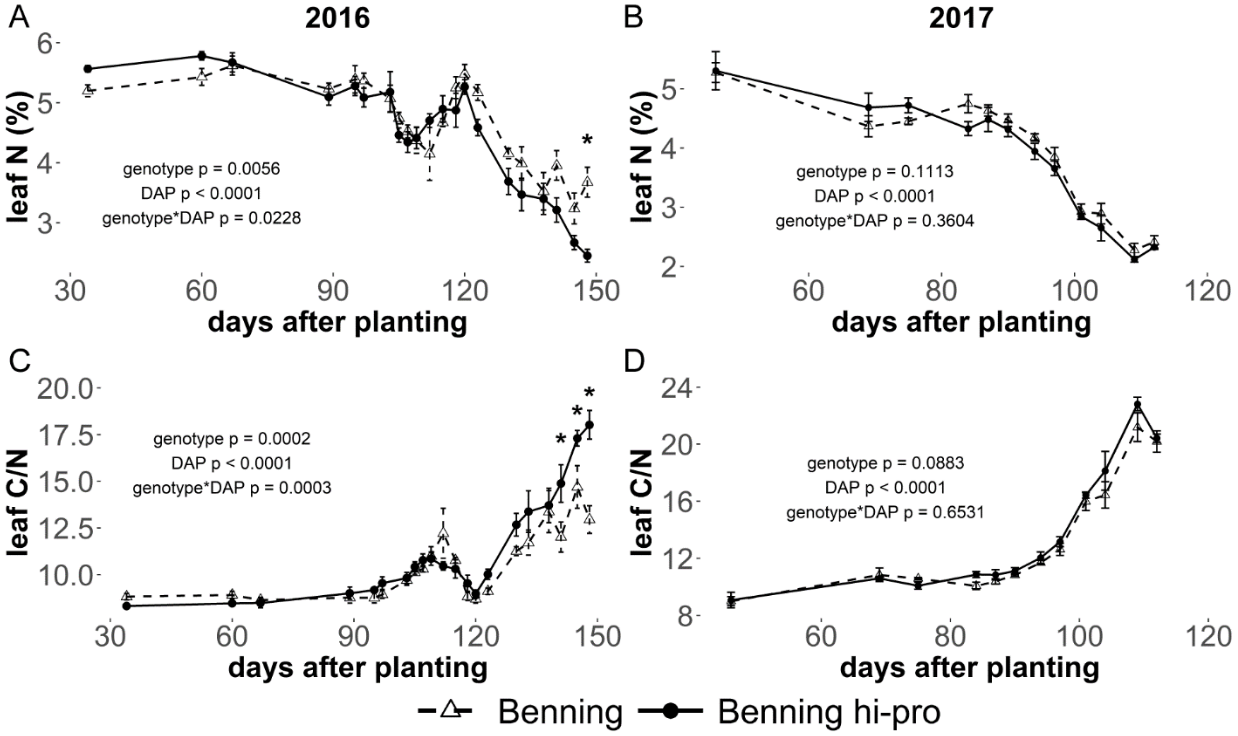

Figure S2. Leaf chlorophyll content during the 2016 seed fill period, R5 and R6. Chlorophyll was isolated from tissue sampled from the uppermost, fully expanded leaf in the canopy. Slopes were not significantly different between the two genotypes.

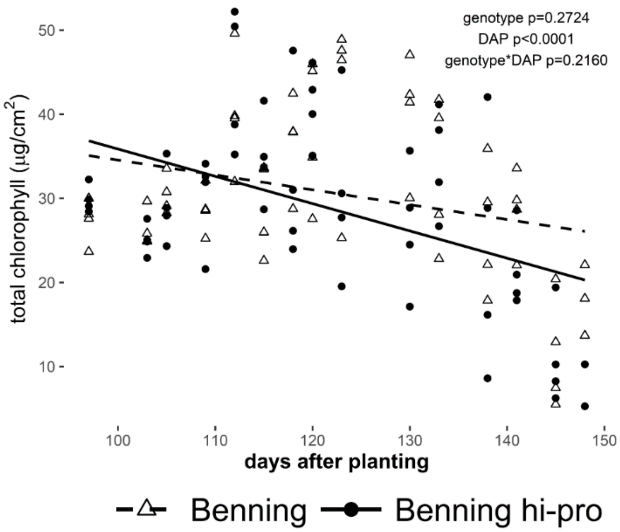
